# Identification of inter- and intra-tumoral molecular phenotypes steering temozolomide resistance in patient-derived glioblastoma cells

**DOI:** 10.1101/2023.08.11.552418

**Authors:** Federica Fabro, Trisha V. Kers, Kate J. Feller, Cecile Beerens, Ioannis Ntafoulis, Ahmed Idbaih, Maite Verreault, Kate Connor, Archita Biswas, Manuela Salvucci, Jochen H.M. Prehn, Annette T. Byrne, Alice C. O’Farrell, Diether Lambrechts, Gonca Dilcan, Francesca Lodi, Ingrid Arijs, Andreas Kremer, Romain Tching Chi Yen, Miao-Ping Chien, Martine L. M. Lamfers, Sieger Leenstra

**Author notes:** **Corresponding Author:** Sieger Leenstra, MD, PhD, Department of Neurosurgery, Brain Tumor Center, Erasmus MC Cancer Institute, Erasmus University Medical Center, Wytemaweg 80, Ee2236, 3015 CN Rotterdam, The Netherlands. Member of the GLIOTRAIN consortium (https://www.gliotrain.eu) (Federica Fabro, Ioannis Ntafoulis, Ahmed Idbaih, Maite Verreault, Kate Connor, Archita Biswas, Manuela Salvucci, Jochen H.M. Prehn, Annette T. Byrne, Alice C. O’Farrell, Diether Lambrechts, Gonca Dilcan, Francesca Lodi, Ingrid Arijs, Andreas Kremer, Romain Tching Chi Yen, Martine L. M. Lamfers, Sieger Leenstra).

## Abstract

**Background:** Radiation therapy and chemotherapy using Temozolomide are the standard adjuvant treatments for patients with glioblastoma. Despite maximal treatment prognosis is still poor largely due to the emergence of Temozolomide resistance. This resistance is closely linked to the widely recognized inter- and intra-tumoral heterogeneity in glioblastoma, although the underlying mechanisms are not yet fully understood. This study aims to investigate the diverse molecular mechanisms involved in temozolomide resistance.

**Methods:** To induce temozolomide resistance, we subjected 21 patient-derived glioblastoma cell cultures to Temozolomide treatment for a period of up to 90 days. Prior to treatment, the cells’ molecular characteristics were analyzed using bulk RNA sequencing. Additionally, we performed single-cell sequencing on four of the cell cultures to track the evolution of temozolomide resistance.

**Results:** The induced temozolomide resistance was associated with two distinct phenotypic behaviors, classified as “adaptive” (ADA) or “non-adaptive” (N-ADA) to temozolomide. The ADA phenotype displayed neurodevelopmental and metabolic gene signatures, whereas the N-ADA phenotype expressed genes related to cell cycle regulation, DNA repair, and protein synthesis. Single-cell RNA sequencing revealed that in ADA cell cultures, one or more subpopulations emerged as dominant in the resistant samples, whereas N-ADA cell cultures remained relatively stable.

**Conclusions:** The adaptability and heterogeneity of glioblastoma cells play pivotal roles in temozolomide treatment and contribute to the tumor’s ability to survive. Depending on the tumor’s adaptability potential, subpopulations with acquired resistance mechanisms may arise. Further research is necessary to deepen our understanding of these mechanisms and develop strategies to overcome them.

## Introduction

The emergence of therapy resistance in glioblastoma still undermines the chance of patients’ survival. Unfortunately, after standard therapy that includes surgery, radiotherapy, and chemotherapy with temozolomide (TMZ), relapses inevitably occur ^1,2^. This is mainly caused by the emergence of therapy resistance, which diminishes the cytotoxic effects of chemotherapy. Understanding the driving molecular characteristics involved in temozolomide resistance has become of utmost importance to improve treatment strategy and patient survival.

Drug resistance is a complex phenomenon that comprises intrinsic and acquired components ^3^. Over the past years, many studies have tried to undercover the processes supporting temozolomide resistance in glioblastoma ^4^. However, most of the *in vitro* studies investigating temozolomide resistance carry some limitations. One means to investigate therapy resistance includes the analysis of the paired primary and recurrent samples. However, this approach is often limited by the availability of the matched recurrent samples, as the benefit of repeated resections upon recurrence is still controversial ^5^. Therefore, many studies rely on the administration of temozolomide to treatment-naïve cell cultures. The suitability of using *in vitro* cell culture as a model for further investigations is supported by the correlation observed between *in vitro* sensitivity to temozolomide and clinical outcome ^6^. However, the exploration of resistance development based on tumor-specific drug concentrations has not yet been comprehensively investigated. In fact, another important source of complexity in glioblastoma derives from its heterogeneous nature, present both among and within tumors ^7^. In the recent years, with the advent of the single cell sequencing, the intra-tumor heterogeneity of glioblastoma has been more deeply characterized, including the identification of specific cellular states that recapitulate four distinct neural cell types ^8^. Since then, several studies have been carried out at single cell level to reveal different aspects of glioblastoma heterogeneity, but only a few have investigated response to drugs ^9-15^. Little is still known regarding glioblastoma heterogeneity in relation to temozolomide resistance and its development over time. Here, to recapitulate the intrinsic and acquired development of temozolomide resistance, we established a panel of 21 temozolomide-resistant patient-derived glioblastoma cell cultures through a prolonged low dose exposure to temozolomide. Given the evidence of the crucial role of heterogeneity in glioblastoma, we explored the inter- and intra-tumoral diversity of temozolomide resistance, to investigate and provide a preliminary insight on the complex phenomenon of resistance and its development.

## Materials and methods

### Cell cultures

Patient tumor tissue was received after routine resections with patient’s informed consent and in accordance with protocols approved by the institutional review board. Glioma stem-like cell (GSC) cultures were established from fresh patient-derived glioblastoma tissue as previously reported ^16,17^. The cultures were tested for mycoplasma infection using the MycoAlert PLUS Mycoplasma Detection Kit (Lonza). The cells were cultured in serum free medium. Further details are described in Supplementary Methods.

### Long-term temozolomide treatment

Each cell culture was kept in medium containing ¼ IC_50_ concentration of temozolomide (Sigma Aldrich) for 90 days. The medium with the drug was refreshed weekly. The treatment started with cell cultures ranging from passages 5 to 9 and extended up to passages 17 to 27.

### Viability assay

To evaluate the sensitivity to temozolomide, cells were tested with a serial 2-or 3-fold drug dilutions (0.0058-3 mM). The viability was measured after 5 days with CellTiter Glo 2.0 (Promega). For more details, see Supplementary Methods.

### Doubling time

Cells were plated in a 96 well plate at seeding density of 1000 cells/well, in triplicate, and incubated at 37°C and 5% CO_2_. Cell counting was performed using a hemocytometer every 24 h for 9 days.

### MGMT promoter methylation

All cell cultures were analyzed to check the methylation status of the MGMT promoter. The DNA extracted from the cells was used to perform a methylation specific PCR. For more details, see Supplementary Methods.

### Bulk RNA sequencing and data analysis

The RNA library preparation was carried out using 2ug of RNA extracted from the cell cultures. The sequencing was performed on an Illumina HiSeq4000. The RNA sequencing data was processed using R. More details on the sequencing and data analysis are described in the Supplementary Methods.

### Single cell RNA sequencing and data analysis

For cell cultures GS359, GS785, and GS789 the cells sequenced comprised five conditions: prior treatment (T0), after the first TMZ exposure when reached the confluency (T1-TMZ and T1-CTRL) and after the last TMZ treatment (T2-TMZ and T2-CTRL). For cell culture GS772, the cell sequenced comprised three conditions: T0, T1-CTRL, and T1-TMZ. Cell cultures were washed with PBS and incubated with Accutase (Invitrogen) until detachment, and collected in HBSS (Gibco, Thermo Fisher Scientific). Subsequently, the cells were centrifuged and resuspended in HBSS and sorted (FACSAria II, BD Biosciences) into 384-well plates based on single cells selection gating. Library preparation was carried out according to the CELseq2 protocol ^18^. Cells were then subjected to single cell transcriptomic sequencing (SORT-seq, ~100k reads/cell). The sequencing and read alignment were performed as described by Muraro et al. ^19^. In total, 4608 cells (each cell culture N=1152) were sequenced. The downstream analysis of the scRNAseq data was performed in R using the Seurat package (v4.0.2) ^20^. More details on data analysis can be found in the Supplementary Methods.

### Validation of single cell RNA sequencing

The markers of temozolomide resistance were validated using the GLASS database as source of clinically relevant recurrent glioblastoma ^21^. Further details are present in the Supplementary methods.

### Immunofluorescence staining

Cells were grown on a glass coverslip until confluence. The immunofluorescence protocol was used for the detection and visualization of the protein of interest (ENO1). See Supplementary Methods for further details on the protocol and analysis.

### Statistical analysis

The half inhibitory concentration (IC_50_) was calculated using GraphPad Prism v.8.4.2 software, using a nonlinear regression analysis. The comparison between controls and treated IC_50_ were calculated in GraphPad applying paired Student’s t-test. The comparisons between non-adaptive and adaptive group were calculated applying the unpaired t-test. A p-values of < 0.05 was considered significant.

## Results

### Glioblastoma cell cultures display two distinct behavioral phenotypes

A total of 21 patient-derived primary wildtype glioblastoma cell cultures were cultured in medium containing ¼ of temozolomide half-maximal inhibitory concentration (IC_50_) for 90 days to induce resistance. The variation of IC_50_ was used as a parameter to assess the acquisition of resistance. After the treatment, the cell cultures displayed two distinct phenotypes. 48% of the cell cultures were characterized by similar IC_50_ when compared to the untreated control, henceforth named non-adaptive (N-ADA) (Figure1A). On the contrary, the remaining 52% of the cell cultures showed a significant increment (2-fold or higher) of the IC_50_ after temozolomide treatment, henceforth named adaptive (ADA) (Figure1A). In addition, the untreated N-ADA cells showed on average a 2-fold higher basal IC_50_ than the untreated ADA cells, suggesting a prevalence of intrinsic resistant populations in the former. As expected from prolonged cell culturing, the IC_50_ values of untreated controls displayed variations over time. Nevertheless, 19/21 cell cultures could consistently be assigned to the same phenotypical category (Supplementary Table 1).

**Figure 1.**
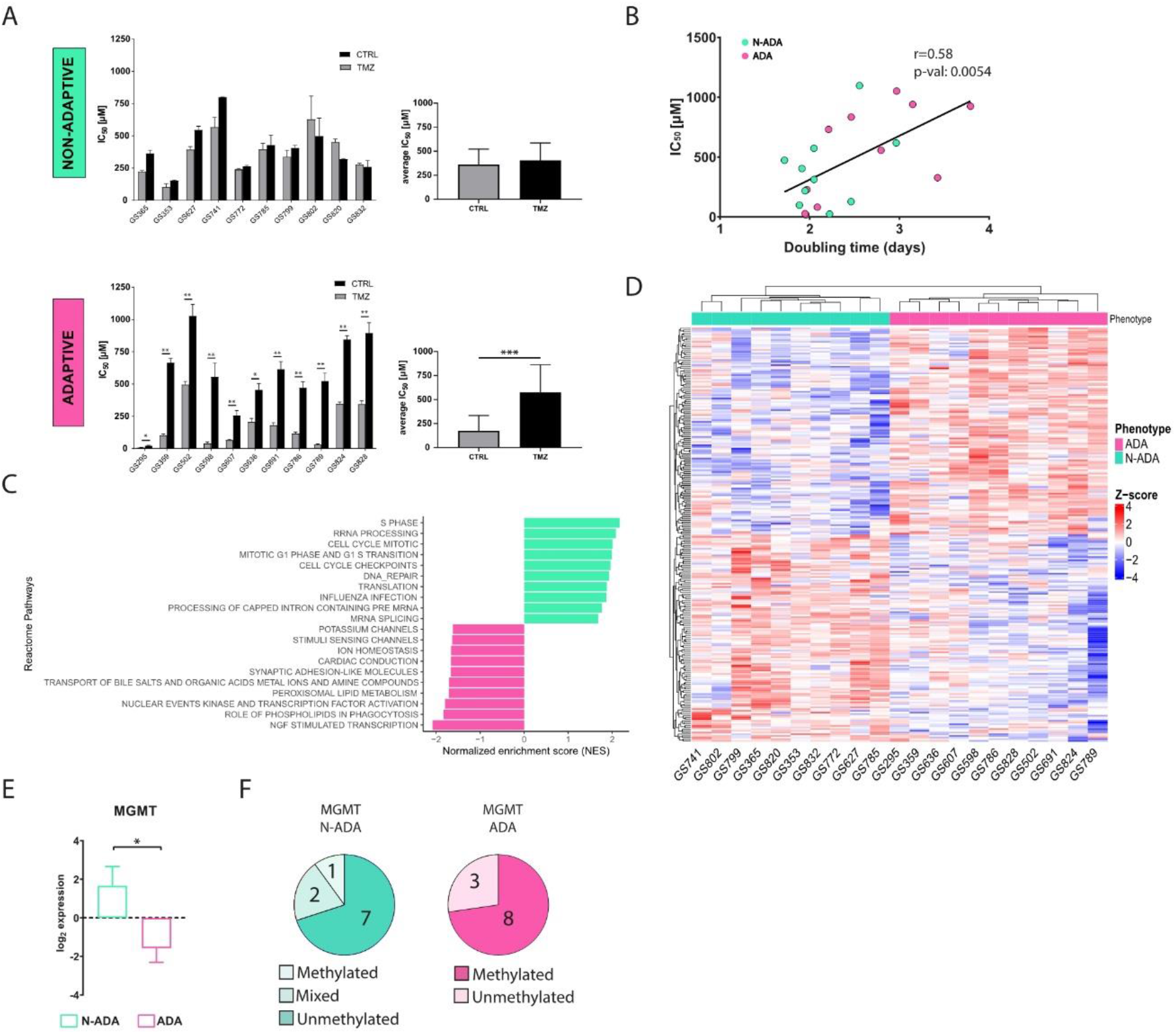
Development of temozolomide resistance in glioblastoma cell cultures. **(A)** IC_50_ values of untreated and temozolomide treated N-ADA (top) and ADA (bottom) cell cultures. On the right, the average of the IC_50_ concentrations. **(B)** Dotplot displaying the correlation between the IC_50_ and doubling times of N-ADA and ADA cell cultures. **(C)** Barplot displaying the top 10 most significant enriched pathways containing the largest number of genes in N-ADA and ADA cell cultures. **(D)** Heatmap of the top 1% most correlated genes for the N-ADA and ADA cell cultures. **(E)** Boxplot showing the gene expression levels of MGMT. The + reports the mean. **(F)** Pie charts showing the amount of N-ADA and ADA cell cultures showing the methylated and unmethylated MGMT promoter status. * p <0.05, ** p <0.01, *** p < 0.001. CTRL: untreated; TMZ: temozolomide. MGMT: O6-methylguanine-DNA methyltransferase.

To characterize the aggressiveness of the cell cultures prior to treatment and its possible relation with the two phenotypes, the growth rate was measured. A moderate positive correlation was observed between doubling time and temozolomide sensitivity of the cells prior to treatment (Figure 1B), indicating a tendency of the less proliferative cells to be more resistant to temozolomide.

### ADA and N-ADA glioblastoma cells express distinct molecular phenotypes prior to treatment

To identify pathways and gene expression signatures underlying the ADA and N-ADA phenotypes, we analyzed the transcriptomic data of the cell cultures prior to treatment by employing gene set enrichment analysis (GSEA). The significantly enriched pathways in the N-ADA group were linked to cell cycle regulation, DNA repair, mRNA and rRNA processing, and translation (Figure 1C). These features have been found to be closely related to aggressiveness and intrinsic drug resistance in glioblastoma ^22,23^. On the other hand, the ADA group was enriched for pathways associated with transcriptional regulation of neuronal system development, lipid-associated pathways, and transport of small molecules. Both neurodevelopment and transcription regulators are known to play a key role in tumorigenesis and plasticity of glioma stem cells (GSC), which are considered the main drivers of glioblastoma heterogeneity ^24^. Metabolic reprogramming is recognized as a hallmark of cancer and a mechanism involved in the adaptation to various stimuli and conditions, requiring among the processes also the regulation of molecular trafficking ^25,26^. Consistently, the top 1% most correlated genes of the N-ADA cell cultures contained genes related to cell cycle regulation and DNA repair (E2F2, FAAP24, CCND1, POLE2, TERT, MGMT), while ADA cell cultures presented genes linked to development (EPAS1, F3, EGR3, SHH, NDN, HAP1) and metabolism including glucose and lipid metabolic genes (DGKG, GPD1, LCAT, ABCD2, ABCA8) (Figure 1D, Supplementary Table 2). Among these genes, MGMT is known to play a crucial role in temozolomide resistance and was significantly upregulated in N-ADA cells (median N-ADA: 3.1; median ADA: -2.6; p-value =0.0157) (Figure 1E) ^4^. Coherently, 70% of the N-ADA and 72% of the ADA cell cultures grouped with the unmethylated and methylated status, respectively, of the MGMT promoter (Figure 1F).

Thus, the development of temozolomide resistance resulted in two distinct phenotypical behaviors prior to treatment suggesting the prevalence of a primary resistance in the N-ADA cells, and an adaptation potential in the ADA cells.

### ADA cell cultures are characterized by subpopulations taking over during temozolomide resistance

In order to examine the temporal changes and heterogeneity of temozolomide resistance within cell cultures displaying increased resistance, we conducted single-cell RNA sequencing at three key stages: prior to treatment (T0), after the first exposure to temozolomide (T1), and following 90 days of temozolomide treatment (T2). This analysis focused on two ADA cell cultures, GS359 and GS789. These cell cultures showed the highest IC_50_ increase over temozolomide treatment following the 90 days of drug treatment. For comparison purposes, we included two N-ADA cell cultures: one encompassing all three time points (GS785) and another covering the first two time points (GS772).

To characterize the composition of the cell cultures, we identified and characterized the main subpopulations present within each sample. Multiple subpopulations were identified and annotated based on glioblastoma-related cell type markers (Supplementary Figure S1A). In addition, we examined the enrichment for Neftel cellular states. Both ADA and N-ADA cell cultures comprised subpopulations exhibiting a mixture of cellular states (Supplementary Figure S1B). The copy number variation (CNV) profile of the tumor cells remained mostly consistent throughout the cell culturing period, as evidenced by the retained CNV profile (Supplementary Figure S1C).

During temozolomide exposure of ADA cell cultures, we observed a progressive increase of one or more subpopulations over time. In GS359, the CSC and OPC subpopulations showed a marked increase in temozolomide-treated cells (CSC: 32%; OPC: 37.3%) compared to untreated controls (CSC: 2.6%; OPC: 7.9%) (Figure 2A). At the point of resistance, the CSC and OPC subpopulations expanded 12 and 4.7 times more, respectively, compared to untreated cells. Similarly, in GS789, the CSC subpopulation showed continuous expansion in temozolomide-treated cells (34.4%) compared to the untreated condition (4.1%) (Figure 2B). Conversely, N-ADA cell cultures exhibited a different behavior. In GS785, although the Astro subpopulation was the most abundant, no difference was observed compared to the untreated condition (Supplementary Figure S3A). The only subpopulation that increased in temozolomide-treated samples was the OPC, with a 2.2-fold increase compared to untreated cells (untreated: 8.7%; treated: 19.4%). In GS772, the subpopulation composition remained unchanged at the beginning of treatment, with the CSC subpopulation consistently being the most abundant (45.2%) (Supplementary Figure S3C).

**Figure 2.**
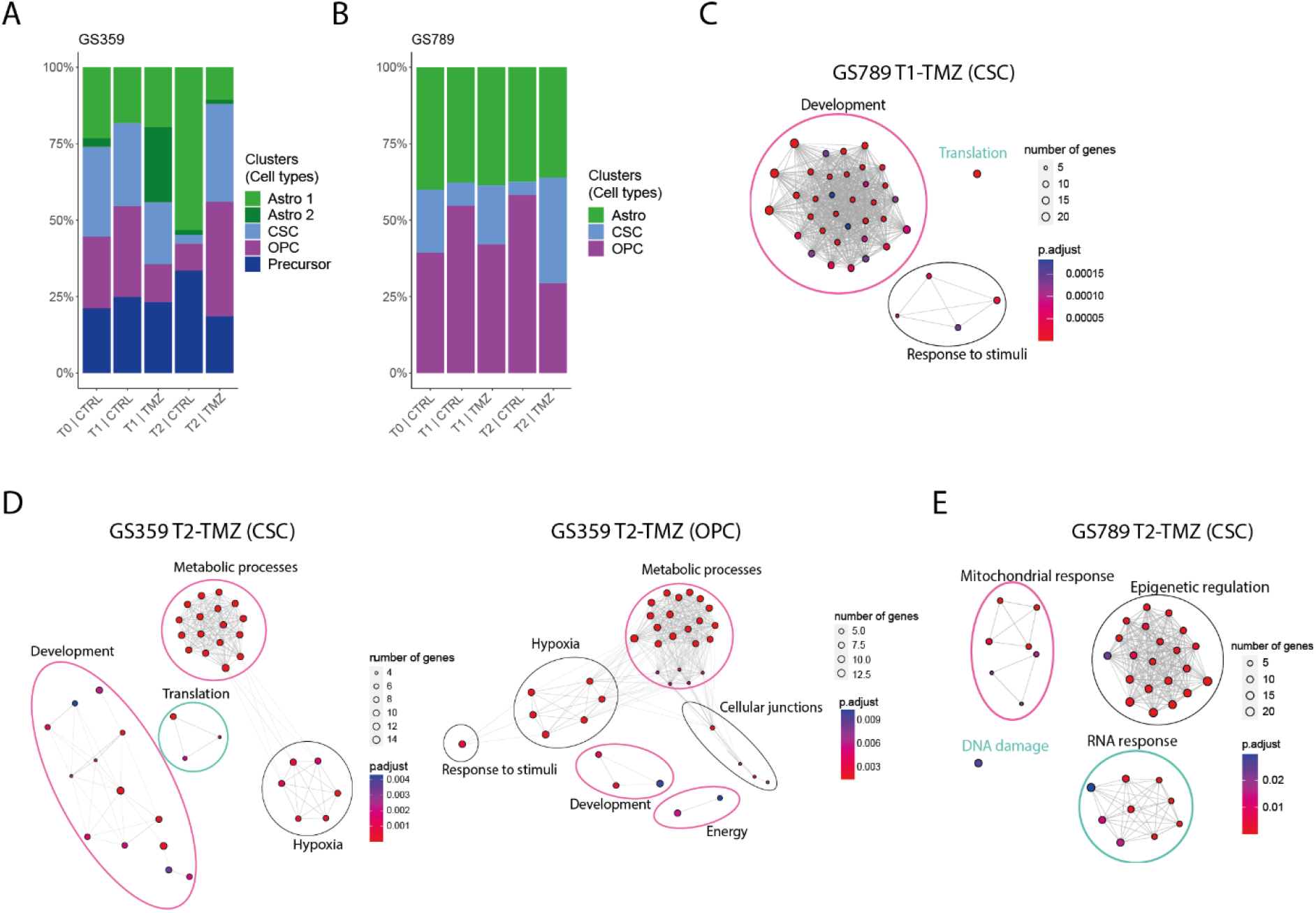
Intra- and inter-tumoral heterogeneity in ADA cell cultures. **(A)** Bar plot displaying the composition of clusters and their relative cell type annotation for GS359. **(B)** Bar plot displaying the composition of clusters and their relative cell type annotation for GS789. **(C)** Network graphs displaying the enriched biological processes identified in T1-TMZ CSC cluster of GS789. **(D)** Network graphs displaying the enriched biological processes identified in T2-TMZ CSC (left) and OPC (right) clusters of GS359. **(E)** Network graphs displaying the enriched biological processes identified in T2-TMZ CSC cluster of GS789. The thickness of the lines in the network graphs represents the percentage of the overlapping genes. Colored ovals identify the major biological processes category related to the clustered processes. Specifically, pink describes ADA-related mechanisms, green represents N-ADA-related mechanisms, and black represents neither ADA nor N-ADA-related mechanisms. CTRL: untreated; TMZ: temozolomide; OPC: oligodendrocyte progenitor-like; Astro: astrocytes-like; Oligo: oligodendrocyte-like; CSC: cancer stem cells.

In summary, ADA cell cultures demonstrated an adaptive behavior, with subpopulations increasing in response to temozolomide treatment, constituting a significant proportion of the tumor cell population. Conversely, N-ADA cell cultures displayed a non-adaptive characteristic, exhibiting less pronounced changes in subpopulation composition during treatment.

### Dominant temozolomide-resistant subpopulations of ADA cell cultures are characterized by distinct resistant biological processes

In order to gain insight into the behavior of the most representative subpopulations in temozolomide-resistant cell cultures during treatment, we conducted an analysis of differentially expressed genes over time (Supplementary Table 3) and their association with biological processes. Using the significantly upregulated genes identified in the dominant temozolomide-resistant subpopulations, we performed gene enrichment analysis to uncover the inter- and intra-tumoral heterogeneity of resistance mechanisms.

At the initial exposure to temozolomide (T1), the CSC subpopulation of GS359 exhibited upregulation of three genes (CHAC1, ARL6IP1, MORN4), while the OPC subpopulation did not show significant upregulation. In GS789, the CSC subpopulation displayed upregulation of genes associated with developmental processes, translation, and response to stimuli (Figure 2C).

Upon the establishment of temozolomide resistance (T2), the subpopulations exhibited more distinct signatures. Both the CSC and OPC subpopulations of GS359 shared genes involved in development, metabolic processes, and hypoxia (Figure 2D). Additionally, subpopulation-specific processes were identified, including translation in the CSC subpopulation, and response to stimuli, cell junctions, and cell energy in the OPC subpopulation. Among the common genes in both subpopulations, ENO1 was identified, a gene associated with metabolic and hypoxia processes known to promote cell growth, migration, and invasion in glioblastoma ^27^. Protein-level analysis revealed increased ENO1 expression over time (fold change: 4.4; p-value < 0.0001) (Supplementary Figure S3A-S3B). Moreover, high expression of ENO1 was correlated with lower overall survival in glioblastomas (Supplementary Figure S3C).

In the CSC subpopulation of GS789, upregulation of genes involved in mitochondrial processes, RNA processes, DNA damage response, and epigenetic regulation, including methylation, was observed (Figure 2E).

Although the N-ADA cell culture GS785 displayed fewer changes in subpopulation composition, temozolomide-specific responses, particularly mitochondrial processes, were still evident in all subpopulations (Supplementary Figure S3B). In GS772, fewer upregulated genes were identified after the initial exposure to temozolomide indicating only a minor initial influence of temozolomide on the tumor cells (Supplementary Figure S3D).

In summary, the results from the dominant subpopulations in ADA cell cultures revealed heterogeneous pathway development involving common genes and biological processes. Despite exhibiting specific responses to temozolomide, both ADA and N-ADA phenotypes appeared to contribute to the development of temozolomide resistance in ADA cell cultures.

### Markers of temozolomide resistance are found in recurrent glioblastoma tumors

To validate the clinical relevance and identify a common pattern in the ADA phenotype, we conducted a study using the GLASS database to investigate the expression of ADA temozolomide resistant markers in recurrent glioblastoma samples.

We considered all subpopulations to determine the overall markers of temozolomide resistance (Supplementary Table 4). As shown in Figure 3A, we found that 37 genes (18 up-regulated and 19 down-regulated) were commonly shared between GS359 and GS789 (Supplementary Table 5). While these genes may not be directly implicated in a particular biological process, the upregulated genes exhibited significant enrichment in cellular components such as collagens, junction complexes, synaptic membranes, filopodia and lamellipodia protrusions, adhesions, and ribosomes (Figure 3B). This implies their potential involvement in essential functions like migration, cell communication, and protein synthesis. Conversely, the downregulated genes were enriched for spindle and endoplasmic reticulum (ER) and vesicle compartments (Figure 3B).

**Figure 3.**
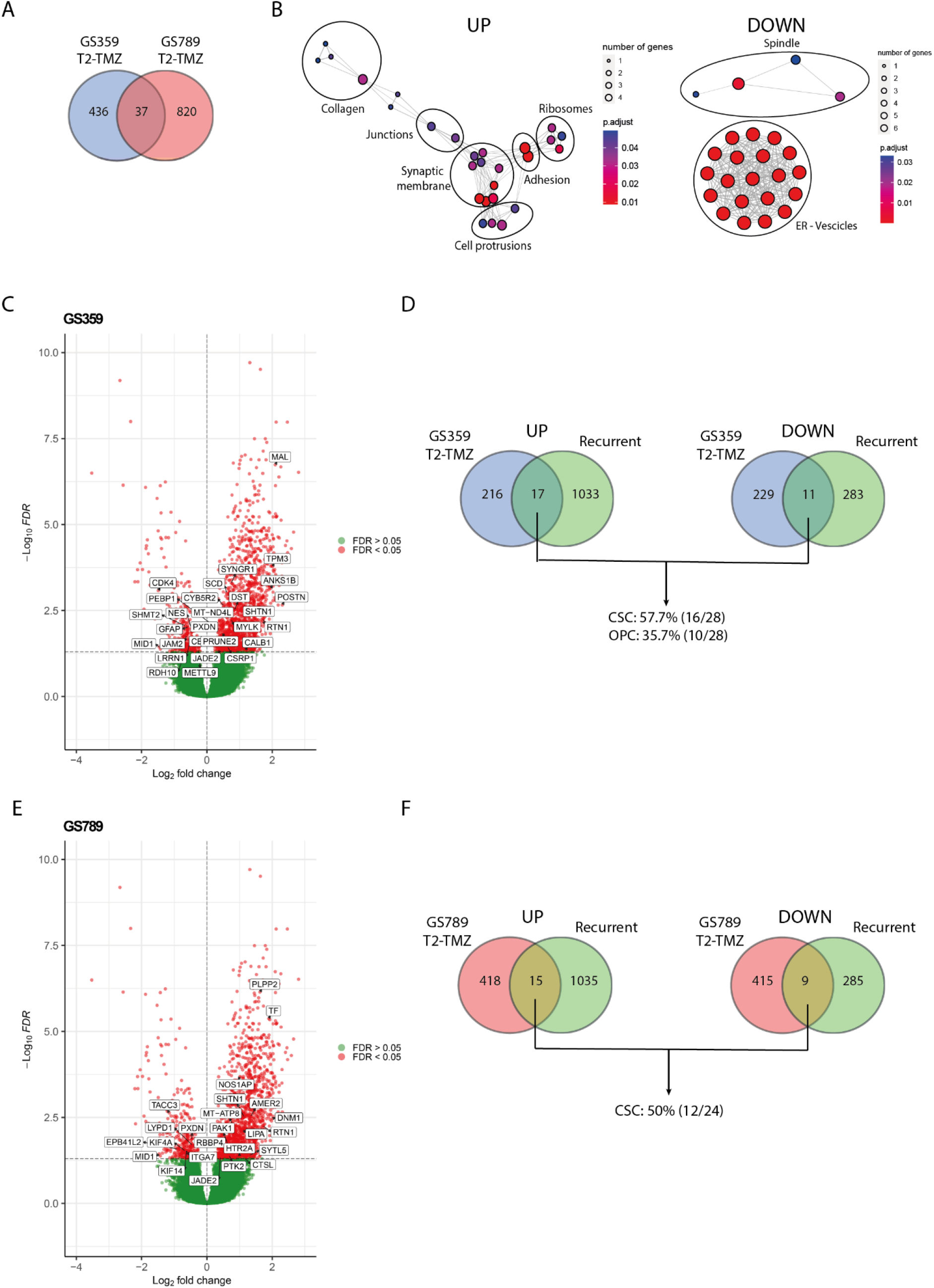
Validation of ADA resistance signature with recurrent glioblastoma. **(A)** Venn diagram displaying the amount of overlapping temozolomide resistant gene signature (T2-TMZ) of GS359 and GS789. **(B)** Network graphs displaying the enriched cellular compartments identified in the common 37 genes of temozolomide resistant (T2-TMZ) ADA cell cultures. On the left, the network graph is related to the upregulated genes, while on the right the network graph refers to the downregulated genes. **(C)** Volcano plot representing the differentially expressed genes between primary and recurrent glioblastomas derived from the GLASS dataset. Positive log2 fold changes refer to the upregulation in the recurrent tumors compared to the primary glioblastomas. The overlapping significant up-and down-regulated genes with temozolomide resistant (T2-TMZ) GS359 are displayed in the white boxes. **(D)** Venn diagrams displaying the amount of overlapping up-(left) and down-(right) regulated genes between recurrent and temozolomide resistant (T2-TMZ) GS359, and the percentage of the genes related to the two dominant clusters. **(E)** Volcano plot representing the differentially expressed genes between primary and recurrent glioblastomas derived from the GLASS dataset. The overlapping significant up-and down-regulated genes with temozolomide resistant (T2-TMZ) GS789 are displayed in the white boxes. **(F)** Venn diagrams displaying the amount of overlapping up-(left) and down-(right) regulated genes between recurrent and temozolomide resistant (T2-TMZ) GS789, and the percentage of the genes related to the dominant cluster. The thickness of the lines in the network graphs represents the percentage of the overlapping genes. Black ovals identify the major cellular compartment category related to the clustered compartments. TMZ: temozolomide.

Furthermore, by comparing the differential gene expression between primary and recurrent wildtype glioblastoma, we identified 3 upregulated genes (SHTN1, RTN1, JADE2) and 2 downregulated genes (PXDN, MID1). SHTN1 is known to regulate neuronal migration and is associated with nervous system development ^28^. RTN1 plays a role in neuroendocrine membrane trafficking, and JADE2 is involved in histone acetylation and chromatin organization ^29,30^. PXDN is a peroxidase involved in the extracellular matrix formation, while MID1 is an E3 ubiquitin ligase implicated in neurodevelopment ^31,32^.

The limited overlap of genes suggests the presence of more tumor-specific patterns associated with resistance and recurrence. To explore this further, we investigated the differentially expressed genes between primary and recurrent glioblastoma in each cell culture (Supplementary Table 5). In GS359, we identified 17 upregulated and 11 downregulated genes (Figure 3C), with 57.7% and 35.7% originating from the dominant temozolomide resistant subpopulations CSC and OPC, respectively (Figure 3D). Similarly, in GS789, we found 15 upregulated and 9 downregulated genes shared among recurrent samples (Figure 3E), with 50% originating from the dominant CSC temozolomide resistant subpopulation (Figure 3F).

We adopted the same approach to examine the T2-TMZ marker genes in N-ADA cell cultures. In comparison to the ADA cell cultures, we observed a similar number of genes detected in recurrent glioblastoma (Supplementary Figure S4A). Lacking a dominant temozolomide-specific subpopulation, we investigated all subpopulations. Interestingly, we observed fewer overlapping genes compared to the ADA dominant subpopulations, with 36.4%, 18.2%, and 0% of the genes across all subpopulations (Supplementary Figure S4B). Likewise, we noted a comparable pattern in GS772 at the initiation of treatment, where the impact of temozolomide is not prominently evident (Supplementary Figure S4C). Overall, the markers of temozolomide resistance of ADA cell cultures that were identified in recurrent glioblastoma specimens reflect the cells’ adaptive response to the drug, indicating the acquisition of new mechanisms to evade its toxicity. These markers exhibit tumor-specific characteristics, underscoring their relevance to individual glioblastomas, and are also enriched for markers associated with the predominant temozolomide resistant subpopulations. The absence of a temozolomide-specific subpopulation in N-ADA cell cultures suggests, instead, a more cooperative response among all tumor cells, in contrast to ADA cell cultures where resistance primarily arise from cellular adaptation of specific subpopulations.

## Discussion

Temozolomide resistance is a persistent issue in glioblastoma treatment. Chemotherapy resistance can be attributed to the heterogeneous nature of glioblastoma, characterized by the presence of multiple subpopulations that do not respond to, or escape therapy ^33^.

The first experimental evidence of this study indicated the presence of two temozolomide resistance behavioral phenotypes. The N-ADA phenotype displayed a phenotypical behavior close to pre-existing resistance phenotypes, where the regulation of cell cycle, DNA repair, RNA processing, and protein synthesis played a major role. Cells use an interconnected network of pathways that regulate cell cycle, DNA repair and replication, as well as transcriptional processes, known as the DNA-damage response (DDR) to preserve the integrity of the genome, leading to cell survival after genotoxic stresses ^34^. These features, in addition to the detection of the unmethylated MGMT in most N-ADA samples, furtherly supported the presence of a main intrinsic resistance signature. In contrast, the ADA cell cultures were mainly characterized by the presence of neurodevelopmental, metabolic, and transport signatures. The main drivers of developmental processes lie in the stem-like population, which has ability and plasticity to drive tumor growth and heterogeneity ^35^. In parallel, the integration of intracellular transport and metabolism becomes a central process of cellular adaptation to environmental changes and cancer progression ^26^. Therefore, the characteristics of ADA cell cultures prior to treatment suggest a crucial role for the ability to adapt to external perturbations, such as temozolomide treatment.

The intra-tumoral heterogeneity observed in the cell cultures appeared to be partially dependent on an adaptive phenotype, as indicated by the increase of specific subpopulations over time in ADA tumors under temozolomide treatment, which was not observed in N-ADA cell cultures. This expansion of subpopulations aligns with the adaptability phenotype of the cell cultures. Interestingly, in both ADA cell cultures, the common expanded subpopulation was the cancer stem cell (CSC) population. Despite the cell culture-specific molecular characteristics, ADA-related mechanisms remained prevalent in ADA subpopulations. The ADA subpopulations were characterized by upregulation of developmental and metabolic genes, which acted as triggers and mediators of resistance. It has also been recognized that developmental signals are responsible for maintaining the CSC population, as CSCs exhibit high metabolic plasticity and can cause therapy resistance through dynamic transitions between distinct metabolic phenotypes ^36,37^. In glioblastoma, different cellular states have been identified along the neurodevelopmental and metabolic axes, which are associated with survival ^38^. Our results additionally suggest that developmental and metabolic features are not only important in sustaining the adaptability and heterogeneity of glioblastoma cells but also in supporting the development of temozolomide resistance in certain subpopulations. Among the markers of resistance, we confirmed the upregulation of ENO1, which is not only a marker of glycolysis but is also involved in other pathological processes such as hypoxia. ENO1 serves as a relevant marker of glioblastoma aggressiveness and poor prognosis ^27^. In the N-ADA cell culture, we did not observe the upregulation of N-ADA specific mechanisms; instead, we observed a variety of different mechanisms, suggesting that their intrinsic characteristics were not utilized and adapted to cope with the drug.

Among the identified subpopulations, we focused on the dominant ones that were most relevant to temozolomide resistance. It is not surprising that the escape mechanisms developed by the resistant subpopulations showed the greatest variability among the cell cultures, each displaying a unique pattern of response to temozolomide. When investigating a common resistance signature among ADA-resistant cell cultures, we identified genes associated with neurodevelopment, particularly those involved in cell protrusions, communication, and extracellular matrix organization. Recent studies have revealed that enrichment of cilium-related genes is characteristic of long-term glioblastoma survivals ^39^. In our study, we propose a new potential role for these genes in governing and regulating temozolomide response, along with cell communication signaling within the tumor environment. When comparing the presence of common temozolomide-resistant gene patterns with recurrent glioblastomas, we did not observe significant overlap. However, when conducting a tumor-specific investigation, more matches were identified, indicating that the development of temozolomide resistance appears to be highly tumor-specific. To validate our findings, we employed the transcriptomic data of recurrent tumor samples in this study. It is crucial to acknowledge that specimens obtained from recurrent patients are highly selected, representing a specific subset of cases as second surgeries are typically reserved for patients who have a more favorable prognosis. Moreover, it is important to consider that tumor tissues vary among cell type populations, as they do not solely consist of tumor cells like this *in vitro* cell culture system. Additionally, recurrent glioblastomas are derived not only from exposure to temozolomide but also from radiation. Finally, the analysis was conducted on a subsample of a more complex tumor model system, and the limited number of tumors analyzed is not sufficient to draw definitive conclusions about the observed trends. Further validations in larger cohorts are necessary. Nevertheless, despite these limitations, our study successfully identified significant genes relevant to temozolomide resistance in clinical samples.

Comprehensive tumor molecular characterization is crucial for identifying drug resistance evolution patterns, as observed in the N-ADA and ADA phenotypes. Regular monitoring is essential to track phenotypic and molecular changes during treatment, aiding treatment decisions. Personalized therapeutic approaches targeting specific resistance pathways in combination with temozolomide may reduce relapse risk. These findings should stimulate further research on drug resistance and heterogeneity in glioblastoma, including microenvironmental factors and identification of molecular targets. Such research holds promise for developing more effective therapies to prevent or overcome resistance in glioblastoma.

## Funding

This project was funded by the European Union’s Horizon 2020 research and innovation program under the Marie Skłodowska-Curie ITN initiative (Grant Agreement #766069).

## Conflict of Interest

The authors declare no competing interest.

## Acknowledgments

The authors gratefully acknowledge the patients who kindly donated tumor tissue and data, making this study possible. In addition, we thank Iris Verploegh for the helpful discussions. M.P.C. acknowledges support from the Oncode Institute, Stichting Ammodo and NWO (Netherlands Organization for Scientific Research) Vidi Grant. M.P.C. appreciates Josephine Nefkens Stichting’s support on the UFO microscope.

## Authorship

Study concept and initiation: SL, FF, and MLML. Acquisition of the *in vitro* data: FF, TK, CB. Acquisition, validation, and preparation of tumor samples: FF, IN, RKB, MV, KC, AB, MS, JHMP. Acquisition of bulk RNA sequencing data: GD, FL, IA, DL. Acquisition of single cell RNA sequencing data: KJF, CB, MPC. Analysis and interpretation of data: FF, MPC, SL, MLML. Obtained funding/project management: AB, AOF, SL, MLML. Drafting of the manuscript: FF. Review and editing: SL, MPC, MLML. Study supervision: SL, MLML.

## Data availability

Bulk RNA sequencing data of the cell lines (n=21) applied in this article is available in the Gene Expression Omnibus Repository: https://www.ncbi.nlm.nih.gov/geo/query/acc.cgi?acc=GSE232173, Accession number: GSE232173. Single cell RNA sequencing data is available in the Gene Expression Omnibus Repository: https://www.ncbi.nlm.nih.gov/geo/query/acc.cgi?acc=GSE226202, Accession number: GSE226202.

## Notes

https://www.ncbi.nlm.nih.gov/geo/query/acc.cgi?acc=GSE232173

https://www.ncbi.nlm.nih.gov/geo/query/acc.cgi?acc=GSE226202

### Competing Interest Statement

The authors have declared no competing interest.

