## Supplementary Figure for "Identification of inter- and intra-tumoral molecular phenotypes steering temozolomide resistance in patient-derived glioblastoma cells"

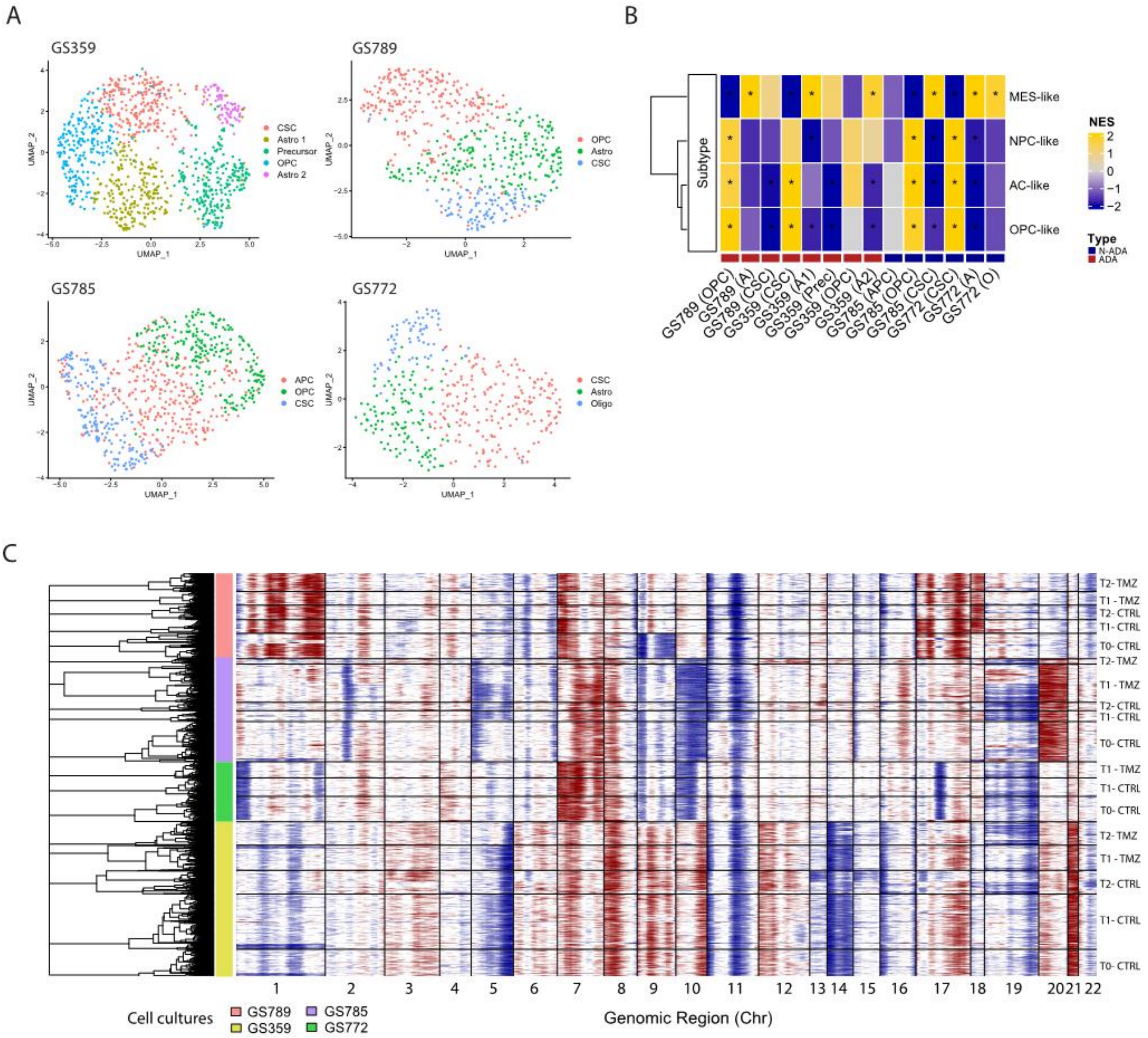


**Supplementary Figure 1. Tumor characteristics at single cell level of ADA and N-ADA cell cultures.**

**(A)** UMAP plot of GS359, GS789, GS785, and GS772 annotated by glioma-related cell types. **(B)** Heatmap of Neftel cellular states subtypes across all clusters and samples. Red and blue bottom legends refer to ADA and N-ADA derivation, respectively. **(C)** Inferred CNV profile of the cell cultures and conditions used in this study. T0-CTRL: initial untreated starting point. T1-TMZ: first temozolomide exposure; T1-CTRL: untreated cells at first temozolomide exposure; T2-TMZ: temozolomide resistant; T2-CTRL: untreated cells at temozolomide resistance. MES: mesenchymal-like; NPC: neural progenitor-like; OPC: oligodendrocyte progenitor-like; AC: astrocyte-like; NES: Normalized Enrichment Score; *: significant. A: Astro; A1/2: Astro cluster 1/2, O: Oligo; Prec: Precursors.

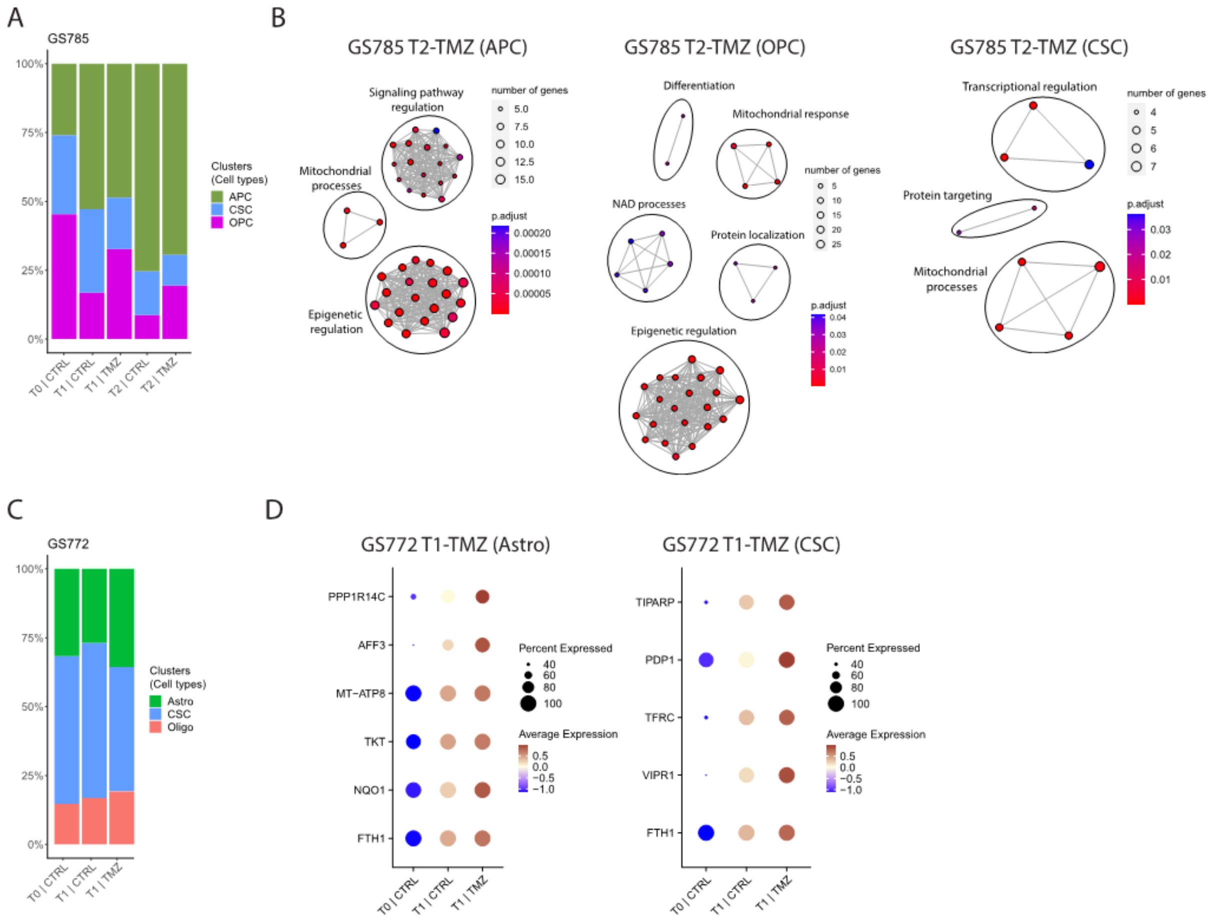


**Supplementary Figure 2. Expression of ENO1 in GS359 ADA cell culture.**

**(A)** Bar plot displaying the composition of clusters and their relative cell type annotation for GS785. **(B)** Network graphs displaying the enriched biological processes identified in T2-TMZ APC (left), OPC (middle) and CSC (right) clusters of GS785. **(C)** Bar plot displaying the composition of clusters and their relative cell type annotation for GS772. **(D)** Dotplot displaying the upregulated genes identified in T1-TMZ Astro (left) and CSC (right) clusters of GS772. None were identified in the Oligo cluster. The thickness of the lines in the network graphs represents the percentage of the overlapping genes. Black ovals identify the major biological processes category related to the clustered processes. CTRL: untreated; TMZ: temozolomide; OPC: oligodendrocyte progenitor-like; APC: astrocyte progenitor-like; Astro: astrocytes-like; Oligo: oligodendrocyte-like; CSC: cancer stem cells.

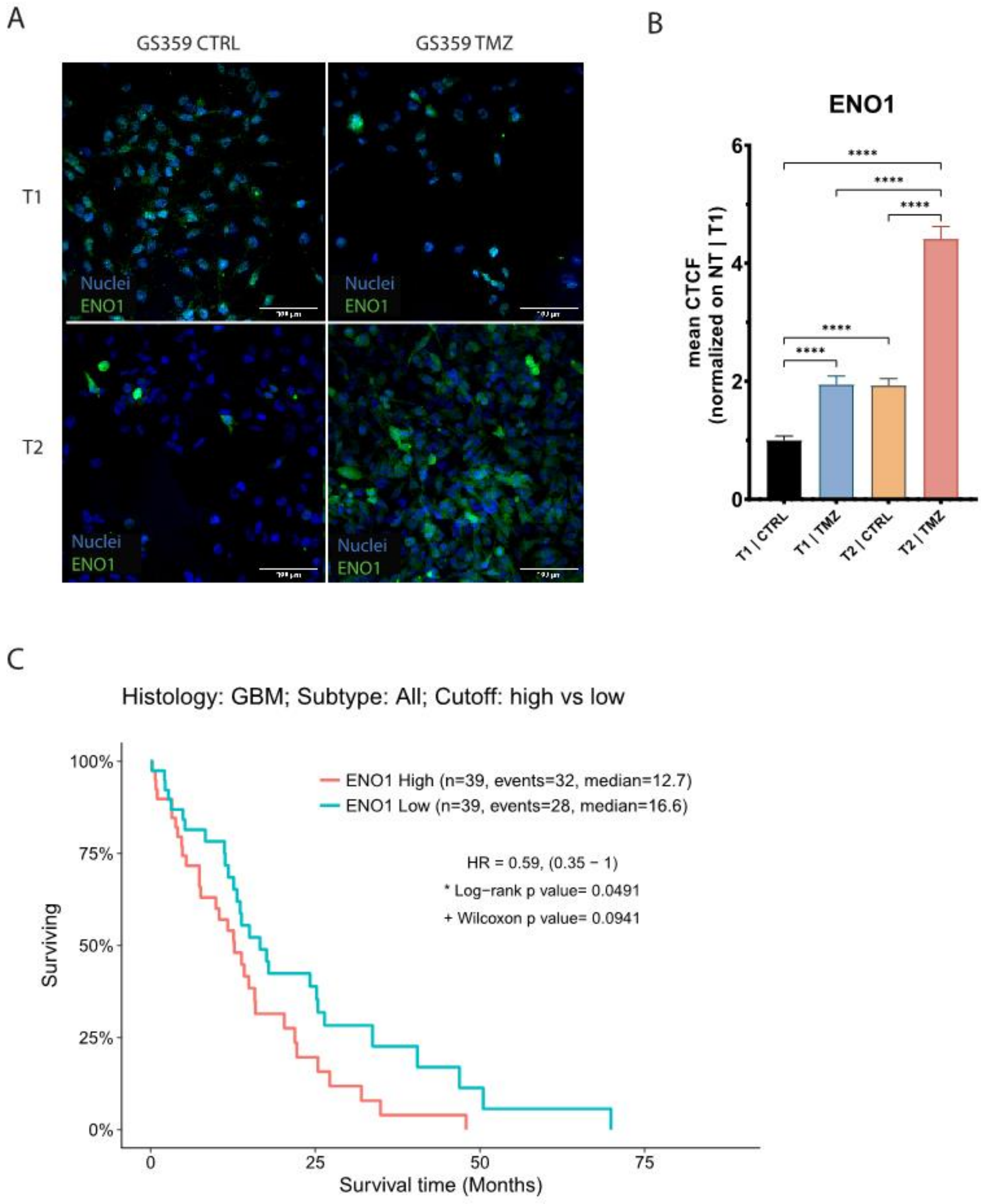


**Supplementary Figure 3. Intra- and inter-tumoral heterogeneity in N-ADA cell cultures.**

**(A)** Representative panel of the immunofluorescence staining of ENO1 in G359 at different conditions (T1-CTRL, T1-TMZ, T2-CTRL, T2-TMZ). **(B)** bar plot represents the normalized amount of ENO1 protein expression during temozolomide treatment. **(C)** Kaplan-Meier graphs showing the overall survival related to high and low expression of ENO1 derived from the TCGA-GBM RNA-seq dataset.

T1-TMZ: first temozolomide exposure; T1-CTRL: untreated cells at first temozolomide exposure; T2-TMZ: temozolomide resistant; T2-CTRL: untreated cells at temozolomide resistance. CTCF: corrected total cell fluorescence.

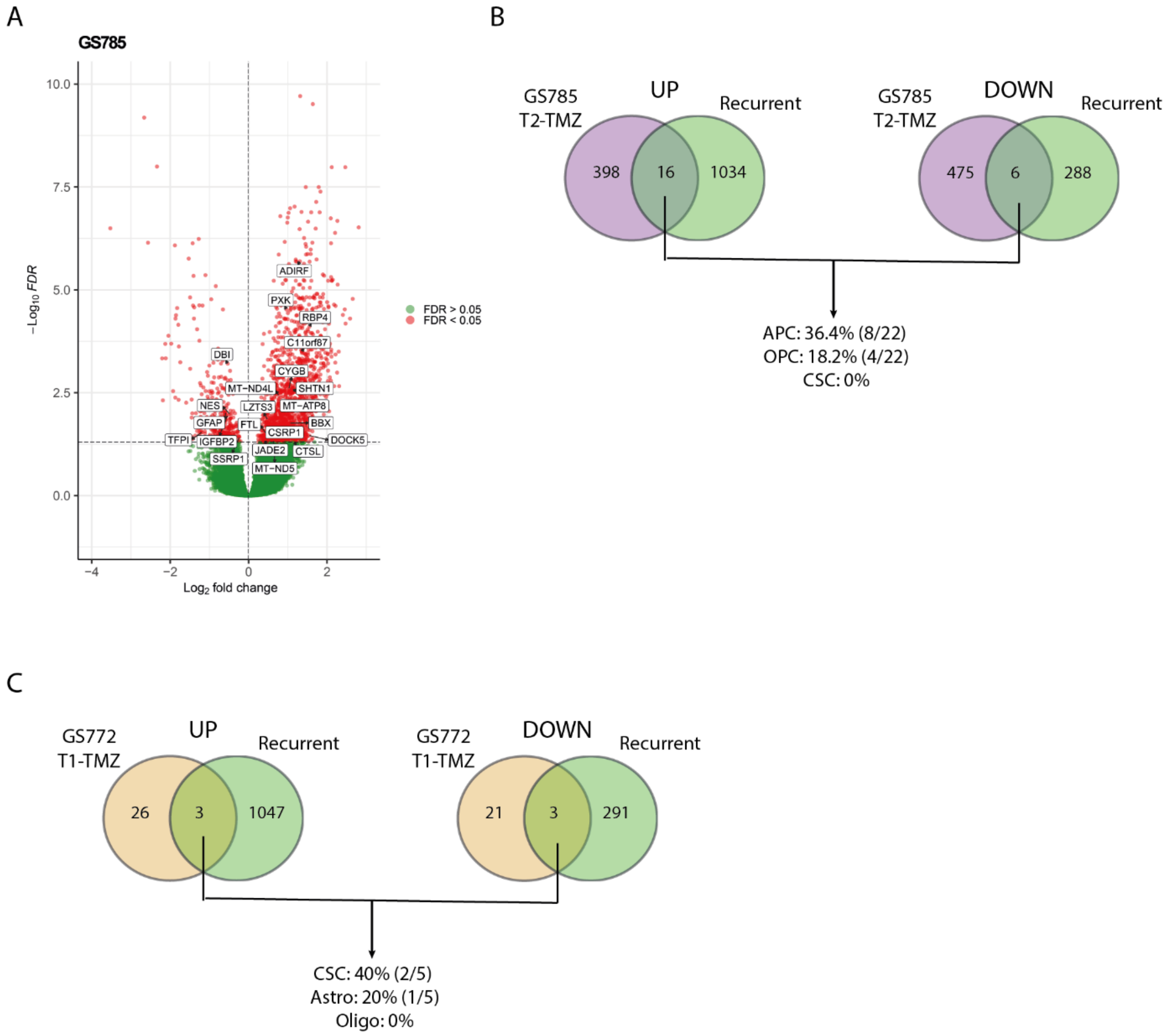


**Supplementary Figure 4**. Validation of N-ADA resistance signature with recurrent GBMs.

**(A)** Volcano plot representing the differentially expressed genes between primary and recurrent GBMs derived from the GLASS dataset. Positive log2 fold changes refer to the upregulation in the recurrent tumors compared to the primary GBMs. The overlapping significant up-and down- regulated genes with temozolomide resistant (T2-TMZ) GS785 are displayed in the white boxes. **(B)** Venn diagrams displaying the amount of overlapping up- (left) and down- (right) regulated genes between recurrent and temozolomide resistant (T2-TMZ) GS785, and the percentage of the genes related to the clusters composing GS785. **(C)** Venn diagrams displaying the amount of overlapping up- (left) and down- (right) regulated genes between recurrent and after the first exposure to temozolomide (T1-TMZ) GS772, and the percentage of the genes related to the clusters composing GS772.
TMZ: temozolomide; OPC: oligodendrocyte progenitor-like; APC: astrocyte progenitor-like; Astro: astrocytes-like; Oligo: oligodendrocyte-like; CSC: cancer stem cells.
