## Supplementary Methods for "Identification of inter- and intra-tumoral molecular phenotypes steering temozolomide resistance in patient-derived glioblastoma cells"

### Cell cultures

Cell cultures were grown in Dulbecco's modified Eagle's medium-F12, supplemented with 1% Penicillin/Streptomycin, 2% B27, 20 ng/mL bFGF, 20 ng/mL EGF (all from Gibco, Thermo Fisher Scientific), and 5 µg/mL Heparin (Alfa Aesar)) in flasks coated with 1:100 diluted Cultrex PathClear Reduced Growth Factor BME (Cultrex, PathClear) [1].

### Viability assay

Temozolomide (Sigma Aldrich) was dissolved in DMSO. The cells were plated in a 96 well plate at seeding density of 1000 cells/well in serum-free medium. The plates were incubated for 24h prior to the drug treatment. After 24h, serial 2-or 3- fold drug dilutions (0.0058-3 mM) were prepared in serum-free culture medium and added to each well in triplicates. The plates were incubated for an additional 5 days. Viability was measured with CellTiter Glo 2.0 (Promega), a luminescent ATP assay, according to manufacturer's instructions. The luminescence was measured with the Tecan Infinite F Plex. The percentage of viability was normalized based on the non-treated control.

### MGMT promoter methylation

All cell cultures were subjected to analysis of the MGMT promoter methylation status. In a separate study, the methylation status of the MGMT promoter in 13 samples was examined [2]. For the remaining 8 cell cultures, DNA extraction involved resuspending the cell pellets in 5% Chelex®100 (Bio-Rad) and adding 10mg/ml of proteinase K (Thermo Fisher Scientific). The Chelex suspension was further heated for 8 minutes at 100°C, vortexed, and centrifuged at 10000g. Following incubation and heating steps, the methylation-specific PCR was conducted using the supernatant obtained as previously mentioned.

### Bulk RNA sequencing and data analysis

Bulk RNA sequencing was performed on 21 primary glioblastoma cell cultures prior to treatment. The RNA library preparation and sequencing for all cell cultures was carried out as previously reported [3]. Briefly, a total of 2 µg of RNA was isolated from the cell cultures, using KAPA stranded mRNA-Seq kit (Kapa Biosystems). The sequencing was single-end 50 bp performed on an Illumina HiSeq4000. Before mapping, the optical duplicates and adaptors were removed with Clumpify v.37.28 and Fastx clipper v.0.0.13, respectively. The quality of the reads was checked with FastQC v.0.11.4. Next, the reads were mapped to the human reference genome GrCh38 with STAR v.2.6 and gene expression matrices were generated with HTSeq v.0.10.0. The raw counts were normalized using the R package EdgeR [4]. The counts were filtered out for the lowly expressed genes and normalized using TMM method in log2 scale. Batch effects were removed using the ComBat-seq package [5].

The enrichment pathway analysis was carried out with the Gene Set Enrichment Analysis (GSEA) v.4.1.0 software, using the Reactome database [6]. The correlation of the genes with the phenotype were ranked by their signal-to-noise ratio. Following the GSEA guidelines, FDR q-values <0.25 were considered significant.

**Single cell RNA sequencing and data analysis**

Briefly, per patient, the low-quality cells were filtered out, followed by data normalization, integration, and cell cycle regression as implemented in Seurat. Finally, the integrated samples were used to identify clusters by UMAP analysis. Differentially expressed genes were accessed through the FindAllMarkers function using the non-parameteric Wilcoxon rank sum test (Bonferroni adjusted p-value < 0.05), to identify markers of drug response using separate and merged clusters, between treatments and time point conditions.

Cells were annotated using scCATCH package, using “Glioma” and “Glioblastoma” as type of human cancer tissue reference [7]. All the genes derived from the differential gene expression analysis identifying the clusters were used to create a ranked list that has been used as the input for the Gene Set Enrichment Analysis (GSEA) v.4.1.0 software [6]. GSEA was run using a module containing the signatures of the GBM cell states identified by Neftel et al. [8].

The differentially expressed genes identifying the drug treatments and time points response having an adjusted p-value <0.05 were used to run the gene enrichment analysis for gene ontology (GO) biological processes (BP) or cellular components (CC). The network GO analysis was performed using the enrichGO and emapplot function of the clusterProfiler package [9].

Gliovis webtool (http://gliovis.bioinfo.cnio.es/) was used to perform the survival analysis, choosing the TCGA-GBM RNA-seq dataset with “high vs low” cut-off parameter [10].

The CNV were inferred using the package inferCNV, using the astrocytes derived from the study GSE171684 as normal cells [11,12].

**Validation with GLASS database**

RNA sequencing transcripts counts derived from the GASS database were downloaded from Synapse (https://www.synapse.org/glass) [13,14]. The samples were selected in order to retain the primary wildtype GBMs and their first recurrence. The preprocessing steps were carried out as described for the current dataset used in this study. Differential gene expression analysis was conducted using the R package EdgeR [4]. The significantly upregulated and downregulated genes were overlapped with the marker genes of temozolomide resistance derived from the single cell analyses to identify the clinical relevant markers.

**Immunofluorescence staining**

At each condition and time point, the cells were grown on glass coverslips. When reached confluence, the cells were washed with PBS and fixed for 15 minutes with 4% PFA (Sigma Aldrich). The cells were then rinsed with PBS three times, permeabilized in 0.3% Triton X-100 (Sigma Aldrich), blocked in 5% goat serum (Abcam) for 1 hour, and incubated with primary rabbit antibody (ENO1, Abcam, 1:200) for 2 hours. The coverslips were then rinsed three times with 3% BSA in PBS. Next, the cells were incubated for 1 hour with an Alexa Fluor 647 F(ab')2-goat anti-rabbit IgG (H+L) antibody (Invitrogen), diluted at 1:1000 as the secondary antibody. The coverslips were rinsed three times with 3% BSA in PBS and mounted on microscope slides using Vectashield hardset antifade mounting medium with DAPI (Vector Laboratories). Fluorescent images of immunolabeled cells were obtained using CX7 confocal microscope (Thermo Fisher) with 20x objective, so that hundreds of cells per sample and per condition were available for analysis.

Images acquired with the CX7 confocal microscope (Thermo Fisher) were analyzed using the ImageJ software v1.53n. The DAPI signal and those from antibodies were used for segmentation of the nuclei and the cytoplasmic compartments, respectively, as well as the boundary of individual cells. After the segmentation, the area, the integrated density, and the mean gray value were measured. For each image, three background areas were used to correct against autofluorescence. For each cell, the corrected total cell fluorescence (CTCF) was calculated as described by McCloy et. al, with the following formula: CTCF = Integrated Density - (Area of selected cell X Mean fluorescence of background readings) [15]. The bar graphs and statistical analysis (One-way ANOVA and Tukey's multiple comparisons test) were performed using GraphPad Prism v.8.4.2.
